# Complement C3aR deletion does not attenuate neurodegeneration in a tauopathy model or alter acute inflammation-induced gene expression changes in the brain

**DOI:** 10.1101/2025.06.16.660026

**Authors:** Yuanyuan Wang, Shristi Pandey, Martin Weber, Man Kin Choy, Tiffany Wu, Tania Chernov-Rogan, Hai Ngu, Oded Foreman, Luke Xie, Jesse E Hanson

## Abstract

Aberrant activation of the classical complement pathway in the brain is implicated in contributing to synapse loss and neurodegeneration in various neurodegenerative conditions. Given that C3aR is a druggable target in the complement pathway, we evaluated the potential of C3aR KO to rescue neurodegeneration in a tauopathy model and neuroinflammatory responses in an acute endotoxemia model. We found that C3aR KO did not rescue tau pathology, microglia activation markers, neurodegeneration, or behavioral abnormalities in tauopathy model mice. While we found that endotoxemia resulted in numerous transcriptional changes including distinct alterations in subpopulations of microglia, astrocytes and oligodendrocytes, C3aR KO did not impact these alterations. Together, our results suggest that the beneficial effects of blocking the complement classical pathway in neurodegeneration models is likely independent of C3aR activation and raise questions about the rationale for therapeutically targeting C3aR for neurodegenerative disease.

## Introduction

The complement classical pathway, which plays physiological roles in developmental synapse pruning ^1^, can also play detrimental roles in the diseased brain and contribute to synapse loss and neurodegeneration in conditions including Alzheimer’s disease and other tauopathies ^2–5^. Classical pathway activation can be initiated by C1q binding to synapses and ultimately results in cleavage of C3 into the C3b fragment which is covalently attached to the synapse surface, and the C3a fragment which is soluble (Fig. 1A). Removal of synapses during development is mediated by microglial recognition of synapses opsonized by C3b via the CR3 receptor ^1,6^. We previously found that C3 knockout (KO) was protective against neurodegeneration in the TauP301S (a.k.a. PS19) model of tauopathy ^7^, and have subsequently found that block of the classical pathway by upstream C1q KO is also protective ^8^. While the benefit of C3 KO in the tauopathy model could be explained via blocking opsonization of neurons by C3b, a contemporaneous study reported that KO of the C3a receptor, C3aR, was sufficient to protect against neurodegeneration in the same mouse model of tauopathy ^9^. Those results suggesting that C3a/C3aR mediates the harmful effects of complement activation were exciting because C3aR is a G protein-coupled receptor (GPCR), a class of protein that is very amenable to inhibition by small molecule drugs, thus raising an attractive avenue for therapeutically targeting the classical pathway in neurodegeneration.

**Fig. 1.**
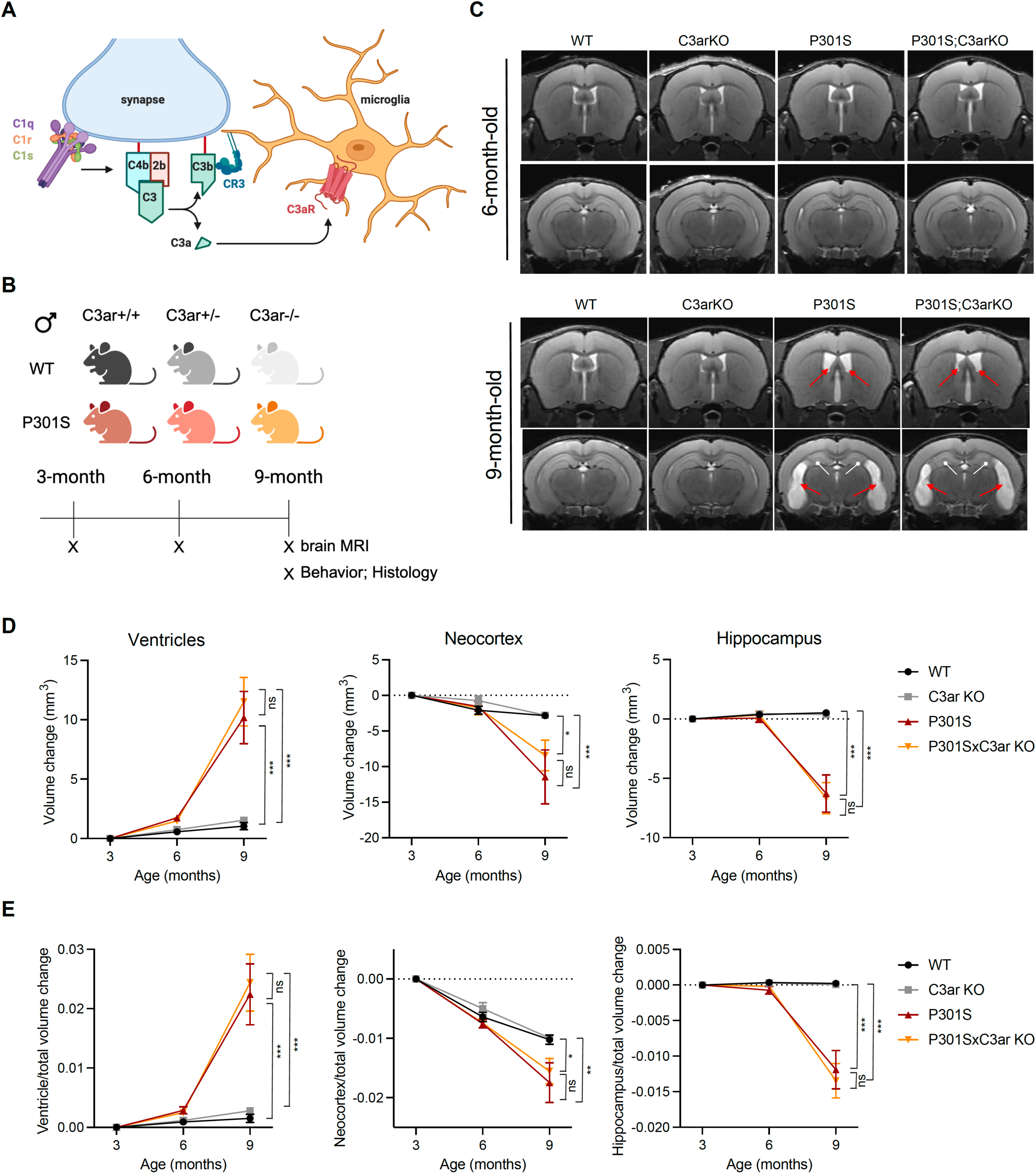
C3aR KO showed no beneficial effects in brain atrophy in TauP301S mice. (**A**) Schematic diagram showing the complement classical pathway function in synapse removal and neuroinflammation via C3b and C3a (created with BioRender.com). C1 complex (C1q, C1r, and C1s) activation and binding to a synapse leads to deposition of C4b and C2b fragments (C3 convertase), resulting in C3b deposition which is recognized by CR3 on microglia, and release of soluble C3a, recognized by C3aR on microglia. (**B**) Study design showing mouse genotypes and timing of experimental measurements: MRI, immunohistochemistry, and behavior. (**C**) Representative MRI images in 6-month-old (top) and 9-month-old (bottom) male mice with genotypes as indicated. Red arrows in 9-month-old TauP301S mouse images indicate enlargement of ventricles, and white arrows indicate hippocampal atrophy. (**D)** Longitudinal MRI quantification of volume changes of ventricles, neocortex, and hippocampus at 6 and 9 months compared with 3 months of age in mice with indicated genotypes. (**E**) Longitudinal MRI quantification of ventricle, neocortex, and hippocampus volumes after first normalizing to total brain volume, showing changes of normalized volumes at 6 and 9 months compared with 3 months mice with indicated genotypes. n = 9-12 male mice per genotype. Data are represented by mean +/- SEM. *, p < 0.05, **, p < 0.01, ***, p < 0.001, two-way ANOVA with Tukey’s multiple comparisons test.

Given the importance of the C3aR KO findings for drug discovery, we aimed to replicate this key experiment and generated C3aR KO mice, crossed them to TauP301S mice, and assessed neurodegeneration, behavior, and pathology. However, using the same assays that showed benefits with C1q KO or C3 KO, we saw no rescue of neurodegeneration or behavior with C3aR KO in the TauP301S mice. We also did not observe any rescue of Tau pathology with C3aR KO. We have previously found that manipulations that alter microglia inflammatory responses, such as loss of TPL2 activity, result in robust rescue of TauP301S neurodegeneration and normalization of bulk brain gene expression changes in response to acute endotoxemia ^10^. Therefore we performed further experiments testing for a role of C3aR in a model of endotoxemia using scRNA-seq analysis of mouse brains following systemic LPS treatment. In these experiments endotoxemia led to numerous changes in the transcriptomic state of different brain cell populations including microglia. This data set provides a previously unavailable analysis including microglia, astrocyte and oligodendrocyte subcluster changes in response to endotoxemia. However, C3aR KO had no effect on any of these neuroinflammatory phenotypes. Overall, we do not find a role for C3aR in acute neuroinflammation or in neurodegeneration in the Tauopathy model, suggesting caution should be taken in considering C3aR as a CNS disease target.

## Results

### C3aR KO mouse generation and validation

To evaluate the potential role of C3aR in neuroinflammation and neurodegeneration, we generated C3aR KO mice using established CRISPR methodology. The successful KO of C3aR was validated by measuring C3aR expression and function (Fig. S1). RT-qPCR from bone marrow derived macrophages (BMDM) showed no expression of C3aR mRNA in the KO mice (Fig. S1A). Flow cytometry analysis of BMDMs from C3aR KO mice stained with a C3aR antibody also showed that C3aR KO mice do not express C3aR protein (Fig. S1B). Moreover, we developed a Ca^2+^ imaging-based assay to examine C3aR function. While the endogenous ligand C3a or the published small molecule agonist of C3aR BR103 ^11^ elicited Ca^2+^ influx in WT BMDMs, the KO BMDMs showed no Ca^2+^ signal in response to the agonists (Fig. S1C), further confirming the loss of C3aR-dependent function in the KO mice.

### C3aR deficiency did not rescue neurodegeneration in Tauopathy mice

Cohorts of littermate mice that were TauP301S positive C3aR +/+ (WT), +/- (HET), and -/- (KO) and TauP301S negative C3aR WT, HET, and KO were all generated from the same breeding pairs. The C3aR HET mice were included in the study for some of the readouts in order to evaluate the therapeutic potential of partial reduction of C3aR function (Fig. 1B). Only male mice were used in this study to avoid confounds of sex differences. To further confirm C3aR knockout in this experimental cohort, we measured C3aR mRNA in all the mouse brains by qPCR, and the expression was as expected (Fig. S1D). P301S mice showed elevated C3aR expression in bulk brain tissue, as is expected given the robust expression of C3aR in microglia and the strong microgliosis in this model^7,10^. At the same time, in both WT and P301S mice we saw complete elimination of C3aR in KO brains and about ∼ 50% C3aR expression in HET brains. Consistent with our previous studies ^2,7,10^, longitudinal MRI showed that TauP301S mice started developing brain atrophy at 6 months of age. At 9 months of age, Tauopathy mice displayed a strong neurodegeneration phenotype, as indicated by significantly increased ventricle volumes and significantly decreased brain volumes, including in the neocortex and hippocampus. In contrast to our previous studies with C3 KO and C1q KO, C3aR KO had no effect on ventricle enlargement or cortical or hippocampal volume decline in TauP301S mice (Fig. 1C,D). Moreover, C3aR deficiency showed no impact on Tauopathy induced body-weight loss or shrinkage of hippocampus area measured from brain sections (Fig. S2).

### C3aR deficiency did not improve Tau pathology or neuroinflammation in Tauopathy mice

Previous studies have shown TauP301S mice develop dramatic Tau pathology marked by accumulation of phospho-Tau (pTau) and microgliosis as the mice age ^7,8,10,12^. We examined Tau pathology using three markers: AT8 (recognizes pTau at Ser202/Thr205), PHF1 (recognizes pTau at Ser396/Ser404) and MC1 (recognizes neurofibrillary tangles) in 9-month-old male mice. C3aR HET or KO did not affect Tau staining in WT mice. TauP301S mice displayed strong Tau pathology compared with WT mice, indicated by all three markers as expected (Fig. 2A,B). In contrast to the previously published study ^9^, C3aR HET or KO had no significant effects on Tau pathology measured by any of these markers (Fig. 2). When we examined microgliosis and microglial activation in Tauopathy mice using IBA1 and CD68 as markers, again we saw no effect of haploinsufficiency or deficiency of C3aR on these neuroinflammation markers (Fig. 2). The phenotype was similar either from whole brain section analysis or hippocampal region analysis. These findings are different from the previous C3aR KO study ^9^, but are consistent with what we previously observed in C3 KO or C1q KO mice, where we saw no change in Tau pathology or markers of neuroinflammation, despite significant amelioration of brain atrophy in TauP301S mice ^7,8^.

**Fig. 2.**
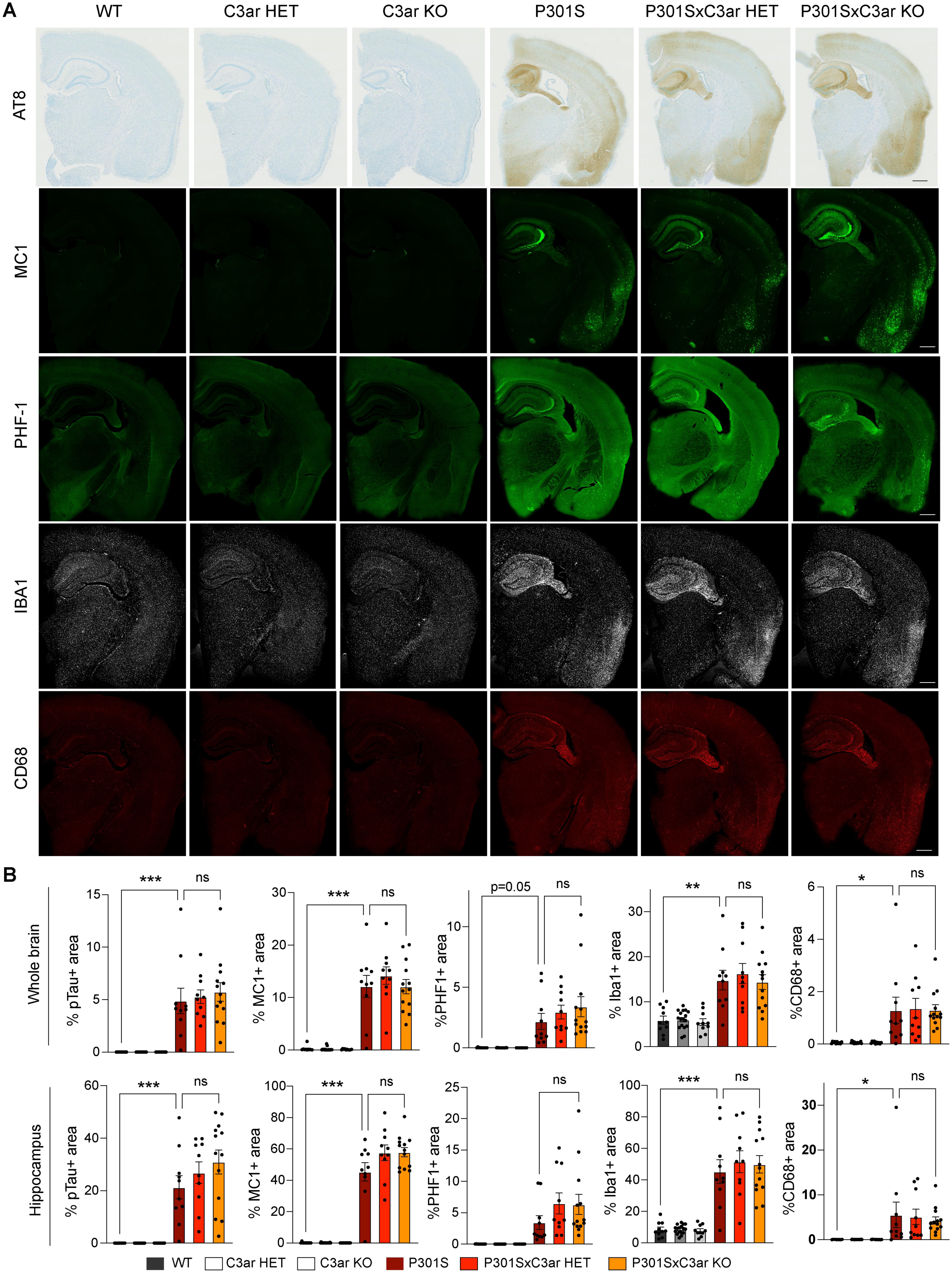
C3aR KO did not improve tau pathology or neuroinflammation markers in TauP301S mice. **(A)** Representative images showing immunostaining for AT8, MC1, PHF-1, IBA1, and CD68 in mouse hemibrains from 9-month-old male mice with genotypes as indicated. Scale bar, 500 mm. (**B)** Quantification of the percentage of AT8 (pTau), MC1, PHF-1, IBA1, and CD68 positive areas within mouse hemibrains (top) and specifically within the hippocampus (bottom) with genotypes as indicated. Data are represented by mean +/- SEM. Each dot represents one animal. n = 10-17 male mice per genotype. *, p < 0.05, *, p < 0.01, ***, p < 0.001, one-way ANOVA with Tukey’s multiple comparisons test.

### C3aR deficiency did not improve behavioral deficits in Tauopathy mice

We next investigated if C3aR HET or KO had any functional benefit at the behavioral level. Consistent with previous studies ^7,10^, in the open field locomotor activity test, 9-month-old male TauP301S mice showed hyperactive behavior compared with non-transgenic mice, as indicated by significantly increased beam breaks within the 15-min time period (Fig. 3). C3aR HET or KO did not affect the locomotor activity in the non-transgenic mice, and did not attenuate the hyperactivity in TauP301S transgenic mice. These results are consistent with the lack of any beneficial effects of C3aR HET/KO on neurodegeneration or neuroinflammation in TauP301S mice.

**Fig. 3.**
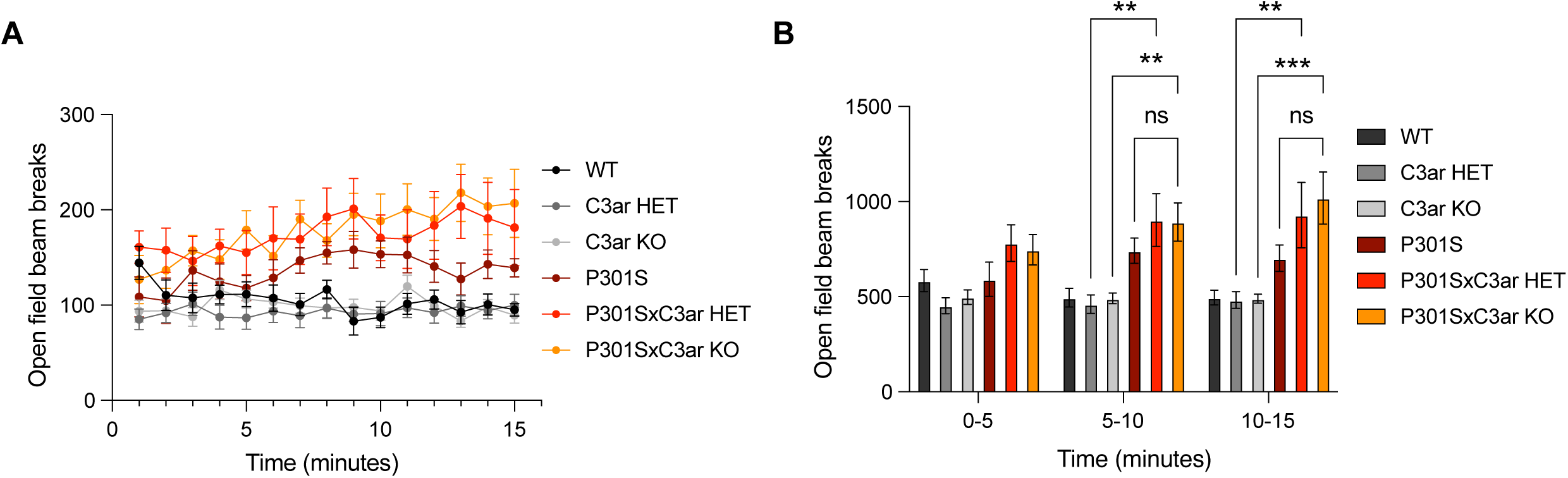
C3aR KO did not attenuate behavioral hyperactivity observed in TauP301S mice. Behavioral spontaneous locomotor activity of 9-month-old mice was evaluated in the open field test by measuring total beam breaks. **(A)** Time course of measured beam breaks by mice with genotypes as indicated. **(B)** Aggregated beam breaks every 5 minutes in mice with genotypes as indicated. Two-way ANOVA with Tukey’s multiple comparisons test, **, p < 0.01, ***, p < 0.001. All data are represented by mean +/- SEM, n = 10-17 male mice per genotype.

### Single-cell RNA-seq analysis reveals glial gene expression changes in response to acute neuroinflammation are C3aR-independent

Since we did not observe any beneficial effects of C3aR HET/KO in the chronic neurodegeneration mouse model, we sought to understand if the C3aR KO has an effect on an acute neuroinflammation model. To this end, we injected mice intraperitoneally with lipopolysaccharide (LPS; 1 mg/kg) or vehicle (PBS) control, and hippocampi were collected after 48 hours. We performed single cell RNA-seq on hippocampi derived from C3aR KO and WT mice that were treated with LPS or PBS. In addition to testing the effects of C3aR genotype, the data also provided an opportunity to investigate glial responses post-LPS treatment in wild type animals, which to our knowledge has not been reported at the scRNA-seq level. After performing standard quality control, dimension reduction, and clustering using Seurat, we recovered 17 broad cell types across the two genotypes and two treatment conditions based on previously defined gene sets ^13–15^ (Fig. 4A, S3A). C3aR expression was predominantly observed in microglia and perivascular macrophages (PVM) (Fig. 4B). Cells derived from LPS-treated animals across both genotypes were also evenly distributed in the tSNE (Fig. 4C). To investigate how LPS treatment and C3aR genotype affected the relative abundance of various cell types, we analyzed the proportion of each cell type across genotype and treatment conditions. No statistically significant differences were observed as a result of LPS treatment or C3aR genotype at the broad cell type level (Fig. S3A). To examine if there are any gene expression changes induced by LPS that are affected by C3aR KO, we performed differential gene expression across the genotypes and treatment conditions in every major cell type identified in our analysis. We found that microglia and PVMs have the highest number of differentially expressed genes following LPS treatment in both genotypes, followed by oligodendrocytes and astrocytes (Fig. 4D).

**Figure 4:**
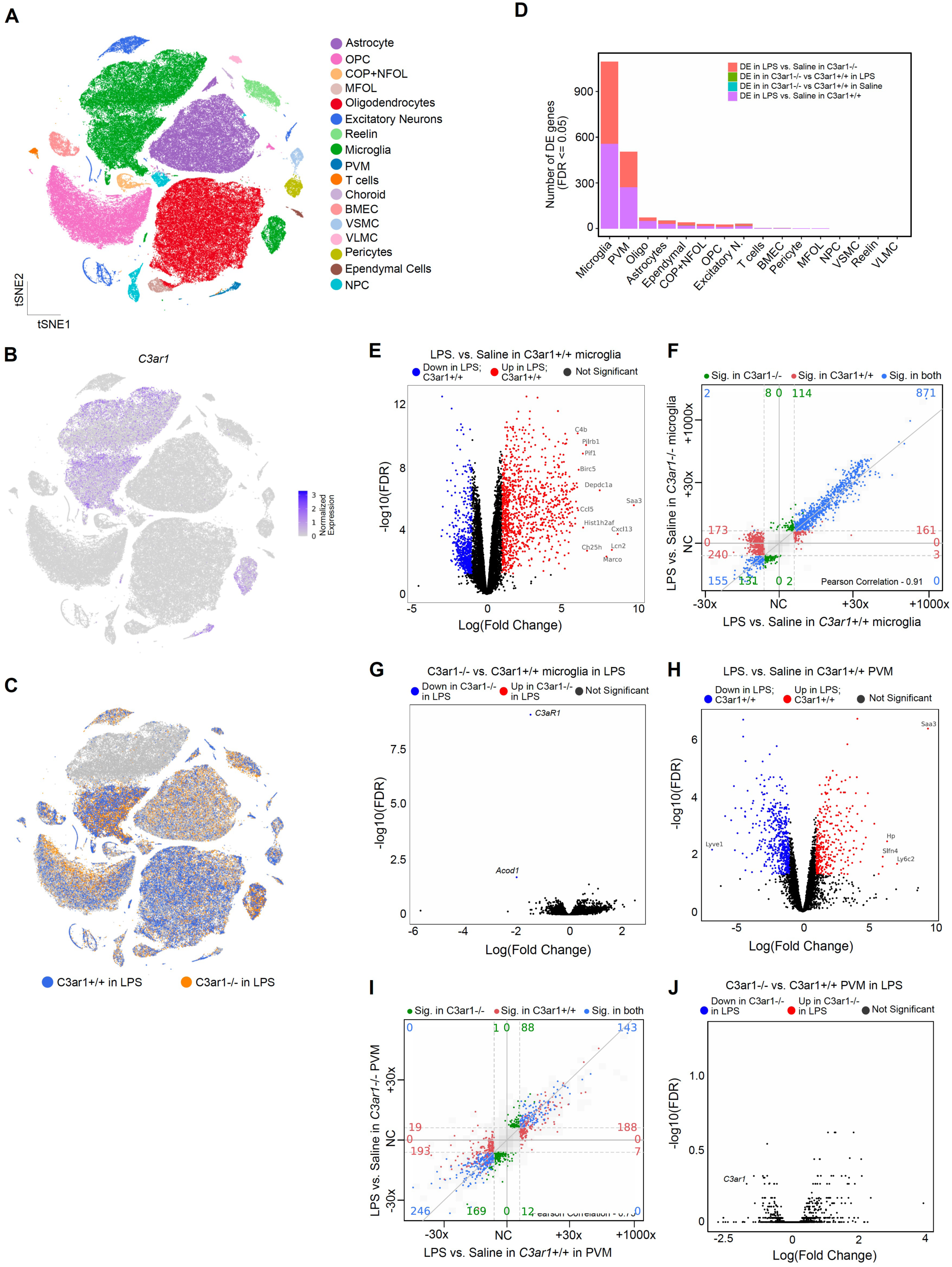
Single-cell RNA sequencing (scRNA-seq) reveals transcriptional changes in glial compartments induced by LPS treatment with no effect of C3aR knockout. Single cell RNA-sequencing was performed on brain tissues from wild-type (C3aR +/+) and C3aR knockout (C3aR -/-) mice treated with lipopolysaccharide (LPS) or vehicle (control). **(A)** UMAP plot displaying the clustering of single cells across both genotypes and treatment conditions, color-coded by the major cell types. **(B)** UMAP plot overlaid with *C3ar1* expression (log-normalized counts), demonstrating that *C3ar1* expression is restricted to microglia and PVMs. **(C)** UMAP plot highlighting the C3aR +/+ and C3aR -/- cells derived from animals treated with LPS. **(D)** Bar plot quantifying the number of differentially expressed genes (DEGs) across all conditions, separated by cell type. **(E)** Volcano plot showing the differential gene expression between LPS-treated and vehicle-treated C3aR WT microglia. Genes with significant changes (FDR < 0.05) are marked with red for upregulated in LPS and blue for downregulated in LPS treatment. **(F)** Four-way comparison of DE genes in LPS treated microglia in C3aR +/+ (x-axis) vs. C3aR -/- genotype (y-axis). Each point represents one gene colored by whether the log Fold change ≥ 2 and FDR ≤ 0.05 in one or both differential expression analysis. (red for C3aR +/+, green for C3aR -/-, and blue for both). Corresponding numbers of DE genes are shown near the borders of the plot. Diagonal line, y = x. **(G)** Volcano plot showing the differential gene expression between C3aR WT and KO microglia in LPS treated mice. Genes with significant changes (FDR < 0.05) are marked with red for upregulated in C3aR KO and blue for downregulated in C3aR KO. **(H)** Volcano plot showing the differential gene expression between LPS-treated and vehicle-treated C3aR KO PVM. Genes with significant changes (FDR < 0.05) are marked with red for upregulated in LPS and blue for downregulated in LPS treatment. **(I)** Four-way comparison of DE genes in LPS treated microglia in C3aR +/+ (x-axis) vs. C3aR -/- genotype (y-axis). Each point represents one gene colored by whether the log Fold change ≥ 2 and FDR ≤ 0.05 in one or both differential expression analysis. (red for C3aR +/+, green for C3aR -/-, and blue for both). Corresponding numbers of DE genes are shown near the borders of the plot. Diagonal line, y = x. **(J)** Volcano plot showing the differential gene expression between C3aR WT and KO PVM in LPS treated mice. Genes with significant changes (FDR < 0.05) are marked with red for upregulated in C3aR KO and blue for downregulated in C3aR KO.

In microglia, which highly express C3aR, we found 556 differentially expressed (DE) genes in LPS vs. control in WT microglia and 540 DE genes in LPS vs. control in C3aR KO microglia at LFC > 2 and FDR <0.05. Genes such as *C4b, Pilrb1, Pif1, Birc5* were upregulated in microglia in response to LPS treatment in WT animals (Fig. 4E). These changes were largely concordant between WT and C3aR KO microglia (Fig. 4F). Notably, aside from C3aR itself, only one gene (*Acod1*) was found to be downregulated in C3aR KO microglia compared to wildtype microglia upon LPS treatment (Fig. 4G).

Next, we investigated the DE genes observed in perivascular macrophages, another cellular compartment with robust expression of C3aR. We found 270 DE genes in LPS vs. saline control in WT and 233 genes in LPS vs. saline control in C3aR KO PVMs (Fig. 4H). Similar to microglia, these changes were largely consistent across the two genotypes and no DE genes wereobserved in the PVMs between C3aR KO and WT animals treated with LPS (Fig. 4I, Fig. 4J). Together, these results suggest that the major gene expression changes induced in microglia in response to LPS treatment are C3aR-independent.

### Microglial cell state changes induced by acute neuroinflammation *in vivo* are C3aR-independent

To further explore gene expression changes induced by LPS and investigate the impact of C3aR on those changes, we subclustered microglia into finer cell states using Seurat’s subclustering. We identified 10 sub-clusters of microglia which were annotated based on marker genes as well as previously defined gene sets (Fig. 5A and B)^16^. As expected, upon LPS treatment, we found that microglia markedly changed their cell states from homeostatic populations to inflammatory cell states, including proliferating and interferon microglia that were described in our previous work ^16^. We found that while the abundance of the homeostatic population significantly decreased in LPS-treated animals, the abundance of the interferon and proliferative microglial clusters, namely, M-C6 to M-C9 were significantly increased in LPS-treated animals. In addition, we also found two LPS-induced populations that we described as LPS1 and LPS2 clusters that were also significantly increased in abundance upon LPS treatment (Fig. 5C). LPS1 was a metabolically active cluster of microglia that was upregulating mitochondrial gene expression and translation and likely transitioning to an activated state, whereas LPS2 represented a fully activated pro-inflammatory microglial state enriched for pathways involved in cytokine production and contributing to inflammatory responses (Fig. 5D, 5E). Both of these LPS-related clusters, in addition to the previously described interferon and proliferative microglial clusters, were induced largely similarly in both WT and C3aR KO hippocampi, showing a statistically significant difference in abundance between LPS and saline control in both genotypes, but no difference in abundance between the two genotypes (Fig. 5C). Together, these results suggest that while microglia proliferate into pro-inflammatory and interferon-related cellular states in acute neuroinflammation, the nature and degree of those cell state shifts were entirely C3aR-independent.

**Figure 5:**
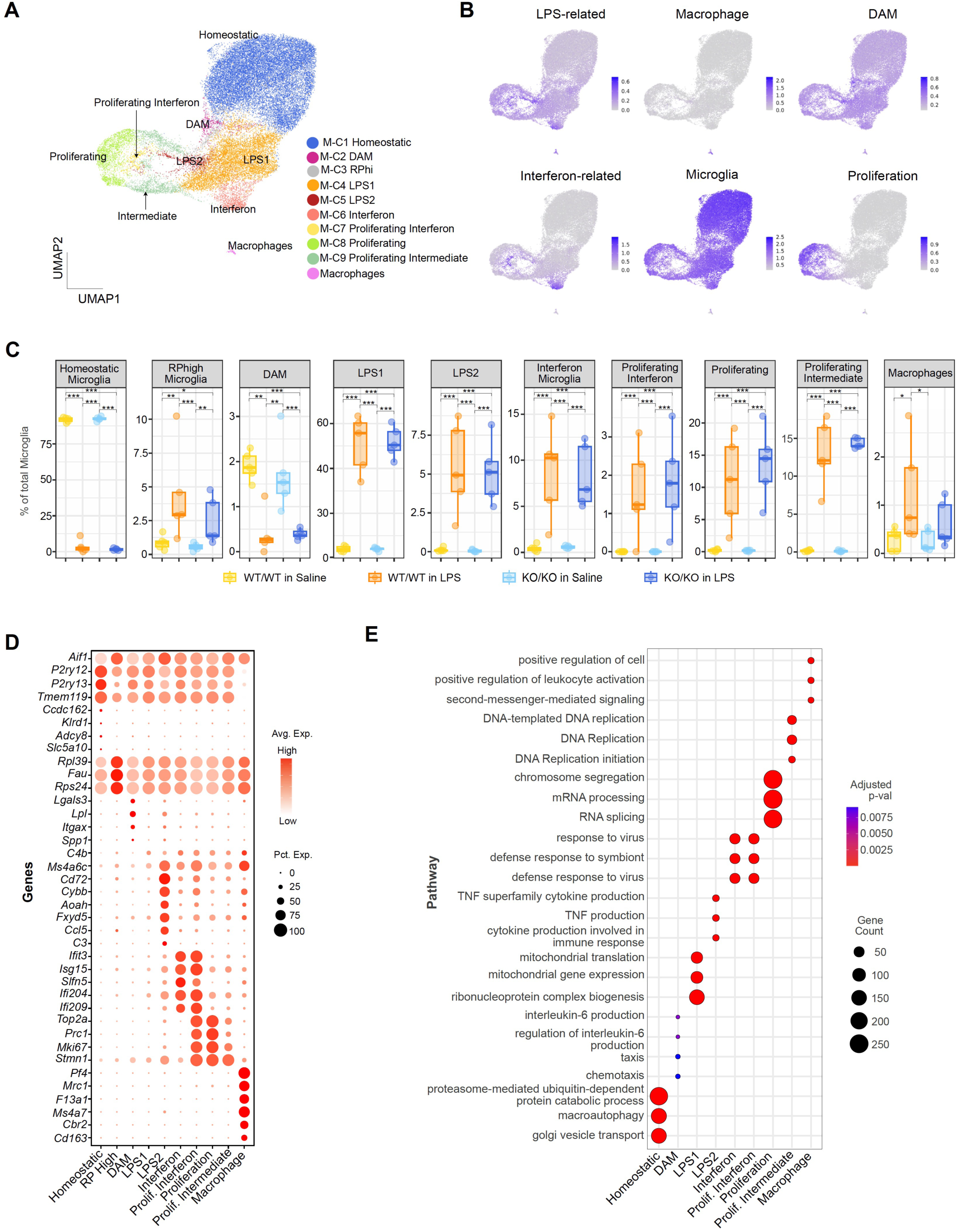
Microglial activation in response to LPS treatment are C3aR independent. **(A)** UMAP plot displaying the sub-clustering of microglia into transcriptionally distinct subpopulations, identified based on single cell RNA-seq analysis. **(B)** Average expression of genes representing specific microglial states calculated across the dataset and represented in the form of a feature plot. **(C)** Proportion of each microglial subpopulation across treatment and genotype groups expressed as a percentage of total microglia, illustrating a shift toward activated microglia in response to LPS treatment, independent of C3aR genotype. Differential abundance statistics * FDR < 0.05; ** FDR <0.01 and *** FDR< 0.001. **(D)** Dotplot of the top marker genes of each microglial subpopulation highlighting the transcriptional states associated with activated and proliferative states. **(E)** Bubble plot representing the results of a Gene Ontology (GO) term enrichment analysis using EnrichGO. X-axis denotes microglial subclusters and y-axis denotes enriched GO terms. The size of the bubble reflects the number of genes contributing to enrichment and the color represents the adjusted p-value.

### Astrocyte gene expression changes induced by acute neuroinflammation *in vivo* are C3aR-independent

In addition to microglia, LPS has also been shown to induce gene expression states in other glial populations ^17,18^. To investigate the effects of LPS on astrocyte states and assess whether knocking out C3aR might have a secondary impact on astrocyte populations, we sub-clustered the astrocytes in our single cell analyses into finer subclusters. Using unsupervised clustering with Seurat, we obtained 7 astrocyte states that were each marked with a unique gene expression profile (Fig. 6A, B). Based on pathway analysis, we found synapse-associated state A-C1, potentially regulatory and adaptive states including A-C5 and A-C2, states potentially involved in structural support such as A-C3, and metabolically adaptive states such as A-C6 (Fig. 6C). All of these states displayed largely similar proportionality across genotype and treatment conditions (Fig. 6D). Finally, we also found an LPS-induced astrocyte state, A-C7, which is marked by high expression of cytokines including interferon stimulated genes such as *Ifit3*, *Ifitm3, Bst2,* and other genes involved in neuroinflammation such as *C4b, B2m* (Fig. 6B). A-C7 is significantly increased in abundance in LPS-treated mice across both genotypes (Fig. 6D) with no effect of C3aR genotype.

**Figure 6:**
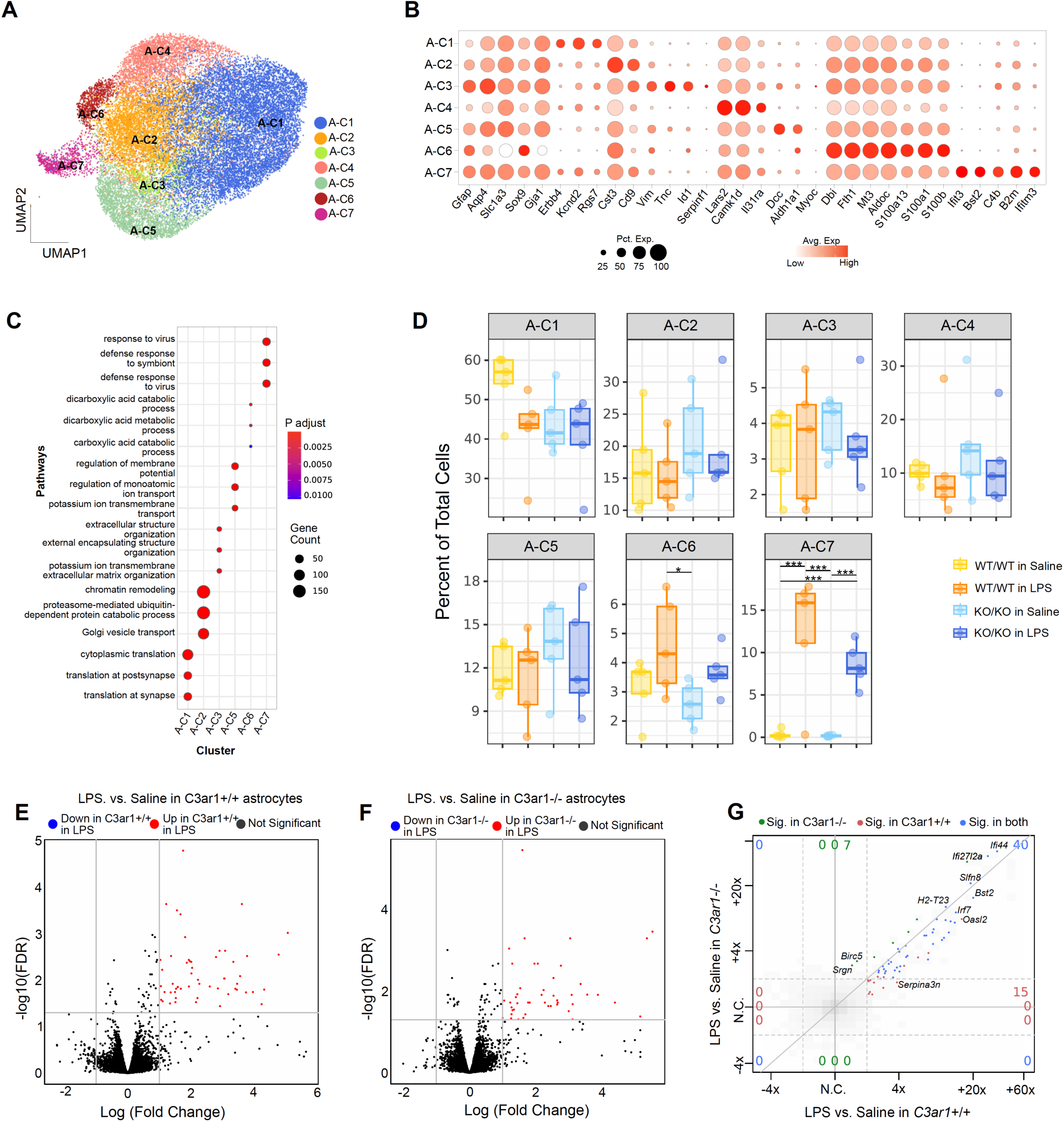
Astrocyte activation in response to LPS treatment are C3aR independent. **(A)** UMAP plot displaying the sub-clustering of astrocytes into transcriptionally distinct subpopulations, identified based on single cell RNA-seq analysis. **(B)** Dotplot of the top marker genes of each astrocyte subpopulation highlighting the transcriptional states associated with LPS treatment. **(C)** Bubble plot representing the results of a Gene Ontology (GO) term enrichment analysis using EnrichGO. X-axis denotes astrocyte subclusters and y-axis denotes enriched GO terms. The size of the bubble reflects the number of genes contributing to enrichment and the color represents the adjusted p-value. **(D)** Proportion of each astrocyte subpopulation across treatment and genotype groups expressed as a percentage of total astrocytes, illustrating a shift toward an inflammatory state in response to LPS treatment, independent of C3aR genotype. Differential abundance statistics * FDR < 0.05; ** FDR <0.01 and *** FDR< 0.001. **(E)** Volcano plot showing the differential gene expression between LPS-treated and vehicle-treated C3aR WT astrocytes. Genes with significant changes (FDR < 0.05) are marked with red for upregulated in LPS and blue for downregulated in LPS treatment. **(F)** Volcano plot showing the differential gene expression between LPS-treated and vehicle-treated C3aR KO astrocytes. Genes with significant changes (FDR < 0.05) are marked with red for upregulated in LPS and blue for downregulated in LPS treatment. **(G)** Four-way comparison of DE genes in LPS treated astrocytes in C3aR +/+ (x-axis) vs. C3aR -/-genotype (y- axis). Each point represents one gene colored by whether the log Fold change ≥ 2 and FDR ≤ 0.05 in one or both differential expression analysis (red for C3aR +/+, green for C3aR -/-, and blue for both). Corresponding numbers of DE genes are shown near the borders of the plot. Diagonal line, y = x.

Examining pseudobulk differential expression in astrocytes (Figure 4D), we found 29 genes upregulated in astrocytes in response to LPS treatment in wild type animals (Fig. 6E) and 24 genes induced by LPS in C3aR KO (Fig. 6F). These genes were largely similar between C3aR WT and C3aR KO (Fig. 6G). Finally, no DE genes were observed between C3aR KO and WT in LPS conditions, suggesting that the effects of LPS on astrocyte gene expression are C3aR-independent.

### Oligodendrocytes gene expression changes induced by acute neuroinflammation *in vivo* are C3aR-independent

Disease-associated oligodendrocytes have been described in various chronic neuroinflammatory contexts ^19,20^. To understand if oligodendrocytes also respond to acute neuroinflammation and to investigate if that response is affected by C3-C3aR signaling, we subsetted the oligodendrocyte lineage cells in our dataset and subclustered them into finer subtypes (Fig. 7A). Using canonical gene sets describing oligodendrocyte subtypes from previous reports ^19,20^, we identified oligodendrocyte precursor cells (OPC), committed oligodendrocyte precursors (COP), newly formed oligodendrocytes (NFOL), myelin forming oligodendrocytes (MFOL), and mature oligodendrocyte (MOL2 and MOL5/6) subtypes (Fig. S4A). Furthermore, we identified a cluster of OPCs (OPC_IFN) that is increased in abundance in animals treated with LPS and characterized by the expression of interferon stimulated genes such as *Ifit1*, *Ifit3* and *Rtp4* (Fig. 7B and 7C). This cluster was also observed in other chronic neurodegeneration models such as TauP301S (Fig S4B). Similarly, we also observed a cluster of oligodendrocytes that is induced by LPS that we named MOL_LPS that bears resemblance to MOL_DA1 and MOL_IFN states described in our previous work (Fig. S4B). MOL_LPS is characterized by increased expression of genes involved in neuroinflammation such as *C4b, Serpina3n,* and *Bst2,* and genes involved in MHC class I antigen presentation such as *B2m, Tapbp, H2-D1,* and *H2-T23.* GO-enrichment analysis highlighted immune response, cytokine production, and immune regulation as top terms enriched in MOL_LPS, similar to previously described MOL_DA1 and MOL_IFN (Fig. 7D). Interestingly, neither of these two LPS induced OPC and OL states were affected by C3aR genotype (Fig. 7B).

**Figure 7:**
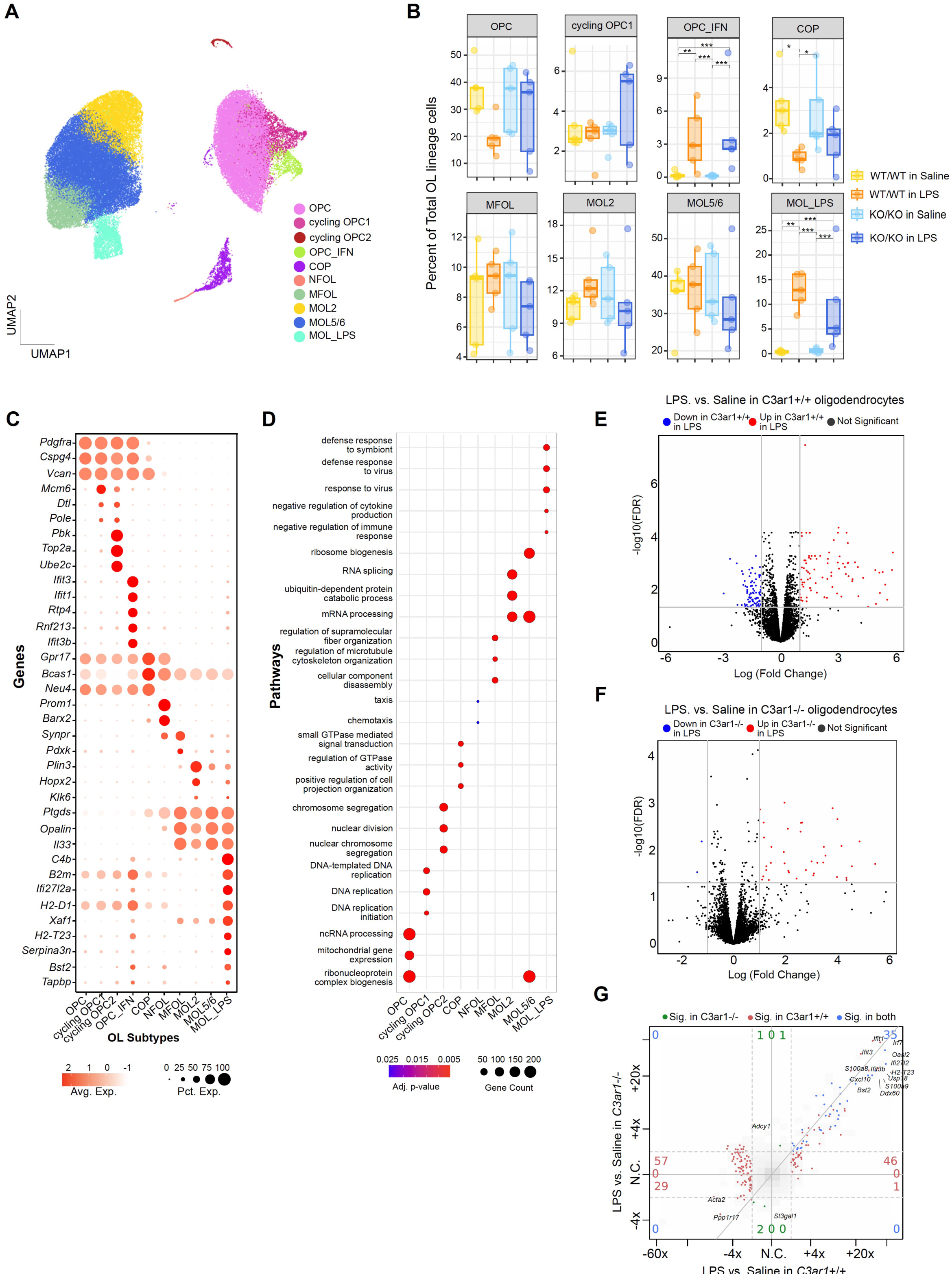
OPC and Oligodendrocyte activation in response to LPS treatment are C3aR independent. **(A)** UMAP plot displaying the sub-clustering of oligodendrocyte lineage cells into transcriptionally distinct subpopulations, identified based on single cell RNA-seq analysis. **(B)** Proportion of each oligodendrocyte lineage subpopulation across treatment and genotype groups expressed as a percentage of total oligodendrocyte lineage cells, illustrating a shift toward an inflammatory state in response to LPS treatment, independent of C3aR genotype. Only cell populations with proportions that are greater than 1% of the total OL lineage cells are shown although all cells were included in the calculation. Differential abundance statistics * FDR < 0.05; ** FDR <0.01 and *** FDR< 0.001. **(C)** Dotplot of the top marker genes of each astrocyte subpopulation highlighting the transcriptional states associated with LPS treatment. **(D)** Bubble plot representing the results of a Gene Ontology (GO) term enrichment analysis using EnrichGO. X-axis denotes astrocyte subclusters and y-axis denotes enriched GO terms. The size of the bubble reflects the number of genes contributing to enrichment and the color represents the adjusted p-value. **(E)** Volcano plot showing the differential gene expression between LPS-treated and vehicle-treated C3arR WT oligodendrocytes. Genes with significant changes (FDR < 0.05) are marked with red for upregulated in LPS and blue for downregulated in LPS treatment **(F)** Volcano plot showing the differential gene expression between LPS-treated and vehicle-treated C3aR KO oligodendrocytes. **(G)** Four-way comparison of DE genes in LPS treated oligodendrocytes in C3aR +/+ (x-axis) vs. C3aR -/- genotype (y-axis). Each point represents one gene colored by whether the log Fold change ≥ 2 and FDR ≤ 0.05 in one or both differential expression analysis (red for C3aR +/+, green for C3aR -/-, and blue for both). Corresponding numbers of DE genes are shown near the borders of the plot. Diagonal line, y = x.

In order to further investigate whether LPS-induced changes in OLs are affected by C3-C3aR signaling, we pseudobulked all the oligodendrocyte clusters by sample and performed differential expression across genotype and treatment conditions. We found 49 DE genes between LPS vs. control comparison in WT oligodendrocytes (Fig. 7E) and 24 DE genes between LPS vs. control comparison in C3aR KO oligodendrocytes (Fig. 7F). The changes observed upon LPS treatment were largely concordant in both comparisons (Fig. 7G) and no DE genes were observed in between KO and WT oligodendrocytes in either vehicle or LPS (data not shown). Together, these results describe oligodendrocyte response in acute neuroinflammation and demonstrate that those responses are C3aR-independent.

## Discussion

We did not observe a role for C3aR in an acute neuroinflammation model or in modulating pathology or neurodegeneration in the tauopathy mouse model. The lack of effect of C3aR KO on tau pathology is consistent with the previously observed lack of a significant effect of KO of the upstream complement components C1q and C3 on pathology in TauP301S mice ^7,8^. Furthermore, using C3aR KO mice, we were unable to see the rescue that we previously observed in TauP301S mice with C1q KO and C3 KO using similar MRI measurements of neurodegeneration and open field measurements of behavioral abnormalities. This is in direct contrast to the benefits previously reported for C3aR KO in this tauopathy model ^9^. Several factors could lead to inconsistent results between studies when using the TauP301S model. Over the course of studying various crosses of TauP301S mice (e.g. C3 KO ^7^, C1q KO ^8^, TPL2 KD KI ^10^, and other unpublished crosses), we have observed that the timing of the onset of pathology and neurodegeneration can vary between cohorts. Therefore, using littermate controls is essential, despite the intensive breeding demands of producing cohorts with all genotypes as littermates. We have also observed that the timing of the onset of neurodegeneration can vary between males and females in some cohorts, therefore it is essential to evaluate males and females independently and avoid mixed cohorts especially with imbalances in sex between the groups. Overall, we have high confidence in our findings that neurodegeneration is rescued in TauP301S mice with C3 KO ^7^ but not C3aR KO (present study), as both experiments were well powered and used single sex littermate cohorts, and all assays and analysis were performed by experimenters who were blind to the genotype of the animals.

It is worth noting that following the original report on C3aR KO in TauP301S mice ^9^, another group reported rescue in TauP301S mice using treatment with the purported C3aR antagonist SB290157 (at 1 mg/kg, 3 times per week for 8 weeks) ^21^. However, there are well established issues with the use of SB290157, including that it can act as a C3aR agonist rather than an antagonist, it exhibits significant off-target activity, and has unfavorable pharmacokinetic properties (making CNS target engagement highly unlikely, especially with this dosing paradigm), which greatly limit the use of SB290157 in probing C3aR function *in vivo* ^22,23^. Furthermore, evidence for CNS target engagement or a pharmacodynamic effect on CNS C3aR function were lacking from that study. Therefore, published experiments with SB290157 are not able to assess the role of C3aR in CNS disease models.

Overall the results of our current and previous studies are consistent with classical pathway activation driving neurodegeneration through C3 cleavage independent of C3aR. This could be mediated by synapse elimination triggered by CR3 recognition of deposited C3b (Fig1A), as has been shown in synapse elimination during visual system development ^6^ and in mouse models of amyloidosis ^4^, virus-induced memory impairment^24^, and Huntington’s disease^25^. In addition, cleavage of C5, which is downstream of C3b, has been proposed to contribute to synapse loss in amyloid models, by either C5aR activation ^26,27^, or formation of the membrane attack complex ^28,29^.

Because we observed no effect of C3aR KO on tau pathology, gliosis, neurodegeneration, or behavior in TauP301S mice, we did not further examine cellular phenotypes in these mice. To check for potential roles of C3aR in an acute neuroinflammation setting, we also examined C3aR KO mice in a model of endotoxemia. This experiment provided a high quality single cell RNA-seq dataset cataloging LPS-induced changes in various subpopulations of microglia, astrocytes, and oligodendrocytes. However, upon knocking out C3aR, we observed no impact of C3aR genotype on LPS-induced gene expression changes in glial cell types or on changes in the cellular proportion of inflammation induced glial states. Overall, the lack of a role for C3aR in acute neuroinflammation or in neurodegeneration in the tauopathy model suggests that the benefits of complement KO in neurodegeneration models are likely independent of C3aR function and that caution should be taken before considering targeting C3aR conditions such as tauopathy. Instead, efforts focusing on other parts of the complement pathway could have greater promise for targeting such conditions.

## Materials and methods

### Animals

Animals were maintained in accordance with the Guide for the Care and Use of Laboratory Animals of the National Institutes of Health. Genentech is an AAALAC-accredited facility, and care and handling procedures of animals were reviewed and approved by the Genentech Institutional Animal Care and Use Committee (IACUC) and followed the National Institutes of Health guidelines. TauP301S mice express human Tau with the P301S mutation, driven by the PrP promoter ^12^, and all experiments in this study used male hemizygous TauP301S mice and non-transgenic littermates as controls. The *C3ar1* KO allele was generated on the *MAPT*^P301S^ transgenic background as described below.

Mice carrying the *C3ar1* KO allele were generated using established CRISPR methodology and electroporation of Cas9 HiFi (IDT) in complex with two synthetic sgRNAs (Synthego): Exon2_5’ 5’-UCAUUUCGCAGACUGAGCCA-3’ ’ and Exon2_3’ 5’-AUGUGUGAAGAUGUGGCCCU-3’) into C57BL/6N zygotes. The KO allele is a 1731 bp deletion spanning *C3ar1* exon2 and part of the 3’UTR. Exon 2 is the only *C3ar1* coding exon. The deleted region corresponds to GRCm38/mm10 chr6:122,849,535-122,851,265. The G0 founder was screened for predicted off-targets as described previously ^30^. Using *in vitro* fertilization (IVF) and oocytes from *MAPT*^P301S^ transgenic females, mice heterozygous for the 1731 bp *C3ar1* KO allele and positive for the *MAPT*^P301S^ transgene were obtained. Further breeding resulted in cohorts with all experimental genotypes produced by the same breedings. . Experimenters were blind to genotype for all behavioral measurements, microscopic, and histological analyses.

### Mouse bone marrow-derived macrophages (BMDMs) culture

Hindlimb bones were extracted from WT or C3aR KO mice. 1-2 bones were placed vertically inside a 500 mL tube with a hole in the bottom, and the tube was placed in a 1.5 mL Eppendorf tube. The pair of tubes were centrifuged for 30s at 5,000g. The pellet in the 1.5 mL tube was resuspended in media containing DMEM, Glutamax, 10%FBS, 1% penicillin/streptomycin, and 50 ng/mL murine CSF-1 (R&D systems, 416-ML-010/CF). Cells were counted and plated in untreated 15 cm petri dishes at 50 million cells/dish. 50% of the media was refreshed with media with CSF-1 every two days. After 5-7 days, the cells were washed with PBS and lifted in PBS with a cell scraper and replated at appropriate densities for downstream assays.

### Ca^2+^ imaging assay

The Ca^2+^ imaging assay was carried out as previously described ^31^. Briefly, mouse BMDMs were plated at 20K/well in a 384-well plate. After 24 hours in the 37°C incubator, cells were incubated for 1hr at 37°C in Ca^2+^ dye loading solution, which contains Ca^2+^ dye, signal enhancer (BD#640178; according to manufacturer’s instructions) and Probenecid (Invitrogen P36400) in assay buffer (HBSS+0.02% BSA+20mM HEPES, pH adjusted to 7.4). Agonists were added and Ca^2+^ responses were measured using FDSS/μCell Image Plate Reader (Hamamatsu) with excitation and emission filters appropriate for the Ca^2+^ dye.

### RT-qPCR analysis of C3aR in mouse BMDMs and mouse brains

For mouse BMDMs, RNA was extracted using the RNAeasy mini kit (Qiagen). Mouse brain tissues were lysed in RLT Buffer (Qiagen) using a TissueLyser (Qiagen). RNA was extracted using the RNeasy Mini QIAcube Kit (74116, QIAGEN) with a QIAcube workstation.

RT-qPCR was performed using qScript XLT One-Step RT-qPCR ToughMix, Low ROXTM (QuantaBio) following the manufacturer’s protocol. RT-qPCR was done on Applied Biosystems ViiA 7 or QuantStudio 7 Flex Real-Time PCR System with C3ar1 gene expression assays (Thermofisher Scientific Mm01184110_m1).

### Flow cytometry

Mouse BMDMs isolated from WT or C3aR KO mice were dislodged using enzyme free PBS based cell dissociation buffer (Gibco) for 20□min at 37°C. Cells were washed and resuspended in cold FACS buffer (1X PBS, 0.5% BSA, 0.05% Na Azide) and kept on ice. Cells were stained with either Alexa Fluor 647-conjugated isotype control antibody (R&D systems, # IC006R; 1:100) or Alexa Fluor 647-conjugated C3aR antibody (R&D systems, # FAB10417R: 1:100) for 30 min at 4°C and then washed with FACS buffer. Sytox blue was added to the FACS buffer at the final step so that live cells could be gated. Flow cytometry was performed using LSRFortessa (BD biosciences), and the fluorescence intensity was analyzed with FlowJo software.

### Longitudinal brain MR imaging

MRI was performed on a 9.4T system (Bruker, Billerica, MA) with a 4-channel receive-only cryogen-cooled surface coil and a volume transmit coil . T2-weighted images were acquired with a multi-spin echo sequence: TR 5.2s, TE 1/spacing/TE12 = 6.5/6.5/78 ms, 56 contiguous axial slices (0.3mm thick), field of view (FOV) = 19.2 mm × 19.2 mm, matrix size 256 × 128, 1 average, and a total scan time of 11 min 6 s. During imaging, anesthesia was maintained with 1.5% isoflurane and rectal temperature was maintained at 37 ± 1°C using a feedback system with warm air (SA Instruments, Stony Brook, NY).

Regional differences in the brain structure were evaluated by registration-based region of interest (ROI) analysis. In brief, each scan was processed and registered to the common coordinate framework v3 (CCFv3, Allen Brain Atlas ^32^ using the ANTx2 processing pipeline ^33^. Multiple echo images were averaged to enhance the contrast-to-noise ratio and corrected for field inhomogeneities. Tissue compartments were segmented using a modified SPMMouse framework. Registration to CCFv3 was performed using ELASTIX, employing a 12-parameter affine transformation followed by non-linear warping. Parametric T2 maps were generated from the multiple echo images and segmented into whole brain tissue and CSF using ANTs ^34^. Regions of interest were extracted from CCFv3 space and corrected using the CSF mask to enhance the accuracy of regional volume estimates.

### Immunohistochemistry and analysis

Mice were deeply anesthetized and transcardially perfused with phosphate-buffered saline (PBS). Hemi-brains were drop-fixed for 48h at 4°C in 4% paraformaldehyde as previously described ^7^. After being cryoprotected and frozen, up to 40 hemi-brains were embedded per block in a solid matrix and sectioned coronally at 30 mm (MultiBrain processing by NeuroScience Associates, NSA) before being mounted onto slides. Brain sections were stained for AT8 at NSA using established protocols (DAB staining). Brightfield slides processed by NSA were imaged on the Nanozoomer whole-slide acquisition system (Hamamatsu Corporation) at 200x magnification. Image analysis was performed using Matlab R2021a (MathWorks) with the image processing, computer vision and deep learning toolboxes. Brain sections were detected using a YOLOv4 convolutional neural network. A semi-automated workflow consisting of a DeepLabv3+ convolutional neural network with a ResNet-101 backbone was used to create the segmentation of the hippocampal region. All hippocampal regions were reviewed by an expert neuropathologist and manually corrected if necessary. The neural networks were trained and validated with Genentech internal historical data and manual ground truth labels. Quantification of chromogenic staining area (AT8) was performed using grayscale and color thresholds ^35^ followed by morphological operations. Positive stain area was normalized to the whole brain section or hippocampal area.

Fluorescent immunostainings were performed using free floating sections. Brain sections were washed with PBS and PBST (PBS with 0.2% Triton X-100), blocked by 10% goat serum in PBST at room temperature for one hour, followed by overnight incubation with primary antibodies in 1% goat serum, PBST at 4°C overnight. After washing, brain sections were incubated in secondary antibodies in 1% goat serum, PBST for 2 hours at room temperature. After PBS washes, brain sections were mounted onto slides with ProLong Gold Antifade Mountant (ThermoFisher Scientific P36931). Primary antibodies used were: MC1 (1:1000) and PHF-1 (1:2000) antibodies were from Dr. Peter Davies, IBA1 antibody (1:1000, Fujifilm Wako, # 019-19741), CD68 antibody (1:1000, Bio-Rad, # MCA 1957). Alexa fluor secondary antibodies were used at 1:500. Immunofluorescent slides were imaged using Olympus VS200 slide scanner and images were analyzed using Qupath software. Data were averaged from three sections per animal.

### Open field Behavior

Spontaneous locomotor activity of 9-month-old males was measured with an automated Photobeam Activity System-Open Field (San Diego Instruments) ^7^. Mice were placed individually in a clear plastic chamber (41L × 41W × 38H cm) surrounded by a locomotor frame and a rearing frame fitted with 16 × 16 infrared (IR) beams to record horizontal locomotor activity (3 cm above the floor) and vertical rearing activity (7.5 cm above the floor), respectively. The total number of beam breaks for both horizontal and vertical movements was measured for a total of 15 min.

### Single cell RNAseq

4-month-old WT or C3aR KO female mice were injected with PBS vehicle control or LPS (1 mg/kg; #L6143, Sigma) (n =5 per condition). 48 hours later, mice were perfused with cold PBS and the hippocampi were immediately sub-dissected. Single cell suspensions were prepared from the hippocampi as described ^10,13^. Briefly, hippocampi were chopped into small pieces and dissociated with enzyme mixes in a Neural Tissue Dissociation Kit (P) (Miltenyi 130-092-628) in the presence of actinomycin D. After dissociation, cells were resuspended in Hibernate A Low Fluorescence medium (Brainbits) containing 5% FBS, with Calcein Violet AM (Thermo Fisher C34858) and propidium iodide (Thermo Fisher P1304MP). Flow cytometry was used to sort and collect live single-cell suspensions for the single-cell RNAseq study. Sample processing and library preparation was carried out using the Chromium Next GEM Automated Single Cell 3’ Library & Gel Bead Kit v3.1 on the Chromium Connect Instrument according to manufacturer’s instructions. Live cell samples were prepared to target 10,000 cells per sample and loaded to the instrument. Libraries were sequenced with HiSeq 4000 (Illumina).

### Single cell data processing, cluster identification, pseudobulk, and differential gene expression

FASTQs were processed with the cellranger pipeline from 10X genomics. Transcript counts for a given gene were based on the number of unique UMIs (up to one mismatch) for reads overlapping exons in sense orientation. Cell barcodes from empty droplets were filtered by requiring a minimum number of detected transcripts. Sample quality was further assessed based on the distribution of per-cell statistics, such as total number of reads, percentage of reads mapping uniquely to the reference genome, percentage of mapped reads overlapping exons, number of detected transcripts (UMIs), number of detected genes, and percentage of mitochondrial transcripts. After this primary analysis step, cells with less than 1,000 total UMIs or greater than 10% mitochondrial UMIs were discarded. Doublets were identified using Doublet Finder and removed from downstream analyses.

Seurat ^36^ was used to calculate PCA, tSNE coordinates and Louvain clustering for all cells (Figures 4A, 5A, 6A, 7A). Cell type markers from ^15,16,19^ were used to identify major cell type clusters, and were interpreted using marker gene sets (Figures S3A). Pseudo-bulk of various cell type expression profiles were derived from single-cell datasets first by aggregating each sample’s data for each cell type as described ^13,14^. So, for n samples and m cell types there were n*m total possible pseudobulks (that is, aggregates of cells of a single type from a single sample). If fewer than 10 cells of a particular type were present in a given sample then they were discarded, so the actual total number of pseudo-bulks was less than n*m. A single “raw count” expression profile was created for each pseudo-bulk simply by adding the total number of UMIs for each gene across all cells of that type from that sample. This gave a gene-by-pseudo-bulk count matrix which was then normalized to a normalizedCount statistic using the estimateSizeFactors function from DESeq2 ^37^, used for calculating gene set scores and visualizing gene expression, and for normalization factors for differential expression (DE) analysis. DE was performed on pseudo-bulk datasets using voom+limma methods for bulk RNA-seq. To put this into more formal notation, let *n_ij_* be the raw UMI number of gene *i* in each cell type *j*. Let *s_j_* indicate the sample of cell *j*. The pseudo-bulk count matrix *B*, with rows indexed by genes and columns indexed by samples (instead of cells) is defined as

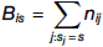

The matrix *B* is then size-factor normalized and analyzed using the standard methods of bulk RNA-seq, including differential expression (DE) analysis using voom+limma ^38^. DE analysis was performed mostly as previously described ^13,14,39^ using voom+limma methods for bulk RNA-seq. For each analysis (that is, for each pair of groups of pseudobulks to be compared), low expressed genes were filtered out. This was defined as genes with at least 10 total UMIs in at least 3 of the analyzed pseudobulks. Gene set scores for each pseudo-bulk profile were calculated as described above for scRNA-seq data, except NormCount values were used in the place of nUMI values.

Cellularity (cellular composition) plots are plotted using <u>geom_boxplot</u> (https://ggplot2.tidyverse.org/reference/geom_boxplot.html). The lower and upper hinges correspond to the first and third quartiles (the 25th and 75th percentiles). The upper whisker extends from the hinge to the largest value no further than 1.5 * IQR from the hinge (where IQR is the inter-quartile range, or distance between the first and third quartiles). The lower whisker extends from the hinge to the smallest value at most 1.5 * IQR of the hinge. Data beyond the end of the whiskers are called “outlying” points and are plotted individually ^40^. FDR values are based on differential abundance testing performed using glmQLFTest function in EdgeR.

Gene set enrichment analysis was conducted using the clusterProfiler package. Marker genes for each cluster were grouped and ranked by descending log2 fold change. The enrichGO function was used to identify enriched Gene Ontology (GO) Biological Process (BP) terms associated with the marker genes. Parameters for the analysis included: OrgDb: org.Mm.eg.db to map mouse gene symbols; KeyType: SYMBOL for gene mapping; Ontology: BP to focus on biological processes; pAdjustMethod: Benjamini-Hochberg (BH) to adjust for multiple testing; qvalueCutoff: 0.05 to filter significant pathways. Only pathways with an adjusted p-value (p.adjust) < 0.05 were considered significant. For each cluster, the top three pathways were selected based on the highest significance (-log10(p.adjust)) for visualization.

### Statistical analysis

Data other than single cell RNAseq data were analyzed using GraphPad Prism. Statistical testing used for each analysis was listed in the corresponding figure legend. Data are represented by mean +/- SEM. Comparisons were considered statistically significant when p < 0.05.

## Resource availability

### Lead contact

Requests for further information and resources should be directed to and will be fulfilled by the lead contact, Jesse E Hanson.

### Materials availability

The mouse strain described in this study is available upon request.

### Data and code availability

Sequencing data have been deposited at GEO (accession number GSE291600) and are publicly available as of the date of publication. Any additional information required to reanalyze the data reported in this paper is available from the lead contact upon request.

## Acknowledgement

We thank our colleagues in the Genetically Engineered Mouse Models (GEM), Microinjection, and Embryo Technology labs for allele design and creation, and our colleagues in the Genetic Analysis Lab and Animal Resources for technical assistance.

## Author contributions

Y.W., S.P., M.W., M.K.C, T.W., T.C., H.N., O.F., L.X., and J.E.H. contributed to the design of the work and interpretation of data. Y.W., S.P., M.W., M.K.C, T.W., T.C., H.N., contributed to the acquisition and analysis of data. J.E.H conceptualized the work. Y.W., S.P. and J.E.H wrote the manuscript. All authors read and edited the manuscript.

## Supplemental information

Supplemental Figures. Fig. S1-S4 and figure legends for Fig. S1-S4.

