## Supplementary Figures (S1-S4) for "Complement C3aR deletion does not attenuate neurodegeneration in a tauopathy model or alter acute inflammation-induced gene expression changes in the brain"

Fig. S1

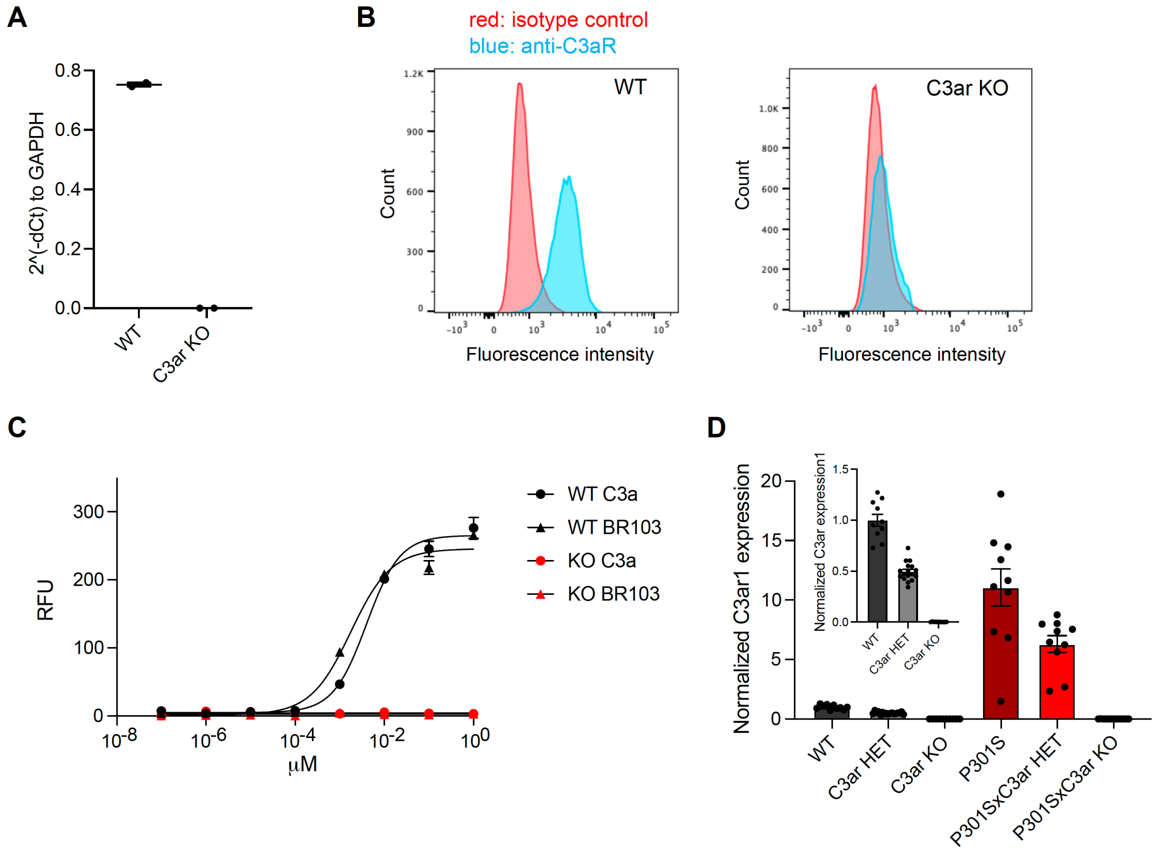


**Fig. S1 Validation of C3aR KO mice.**

**(A)** Expression of *C3ar1* mRNA in bone marrow derived macrophages (BMDMs) isolated from WT or C3aR KO mice as indicated, measured by RT-qPCR after normalization to GAPDH expression in the same samples. **(B)** Live staining of BMDMs isolated from WT or C3aR KO mice as indicated with C3aR antibody (blue) or isotype control antibody (red). Data are shown in histograms of fluorescence intensity measured by flow cytometry. **(C)** Dose response curves of $Ca^{2+}$ signal in response to C3aR agonists C3a or BR103 in WT (black) or C3aR KO (red) BMDMs. WT C3a, EC50 = 0.0040 μM; WT BR103, EC50 = 0.0018 μM. **(D)** Relative expression of *C3ar1* mRNA in the brains of mice with genotypes as indicated, measured by RT-qPCR. *C3ar1* expression was normalized to the average mRNA level in WT mouse brains after normalization to GAPDH in the same samples. Inset shows *C3ar1* expression in TauP301S negative mice with zoomed in y-axis values. Data are represented by mean +/- SEM, n = 10-17.

Fig. S2

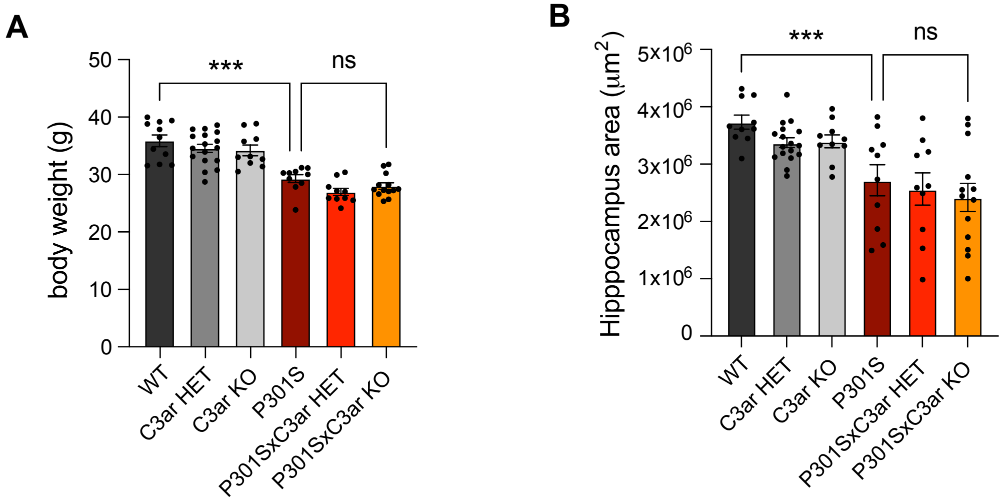


**Fig. S2 C3aR KO did not rescue bodyweight loss or shrinkage of hippocampal area in TauP301S mice.**

**(A)** Body weights of 9-month-old male mice with genotypes as indicated. **(B)** Hippocampal area measured from hemibrain sections of 9-month-old male mice with genotypes as indicated. Data are represented by mean +/- SEM. Each dot represents one animal. n = 10-17 male mice per genotype. ***, $p < 0.001$, one-way ANOVA with Tukey's multiple comparisons test.

Fig. S3

A

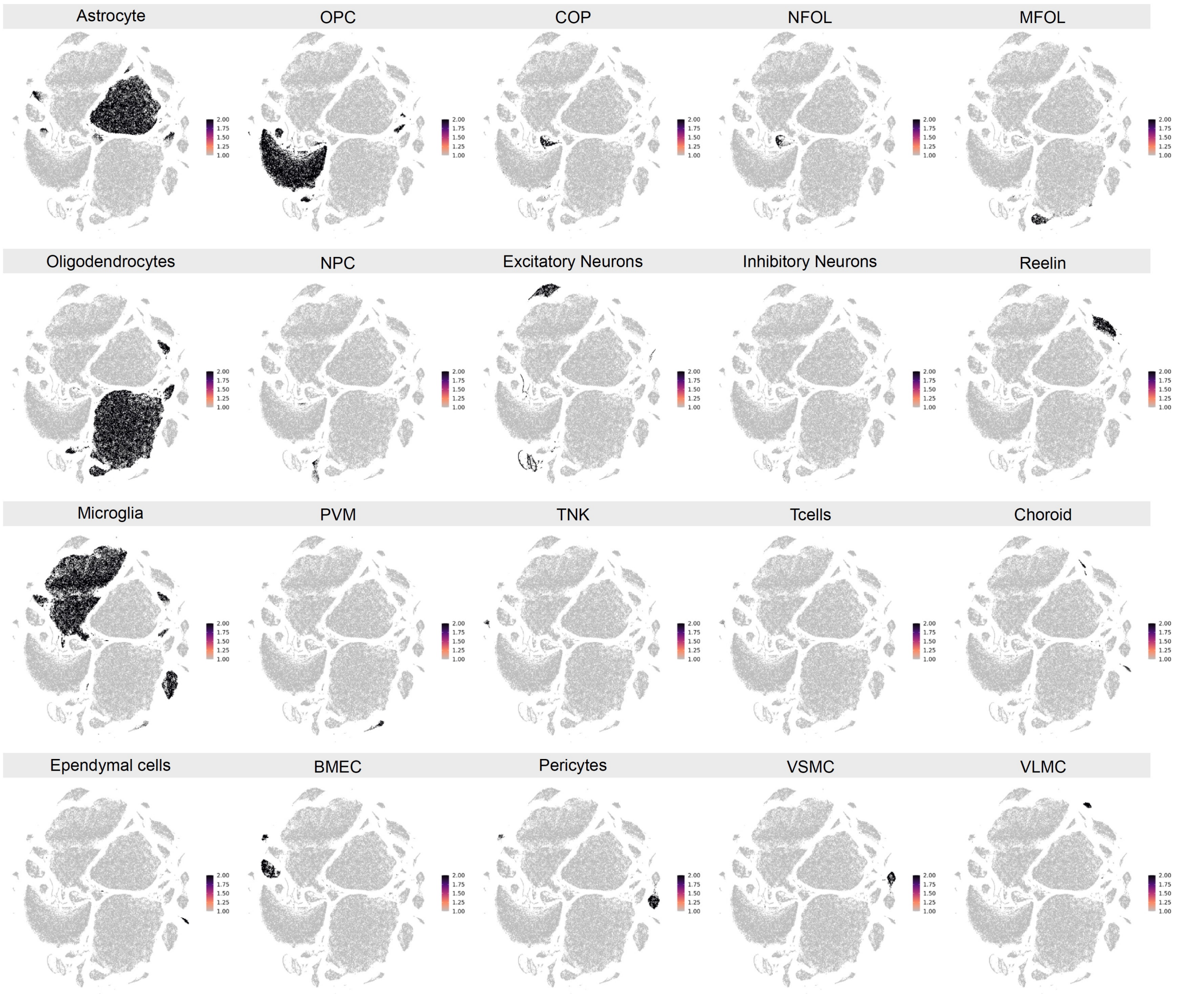


B

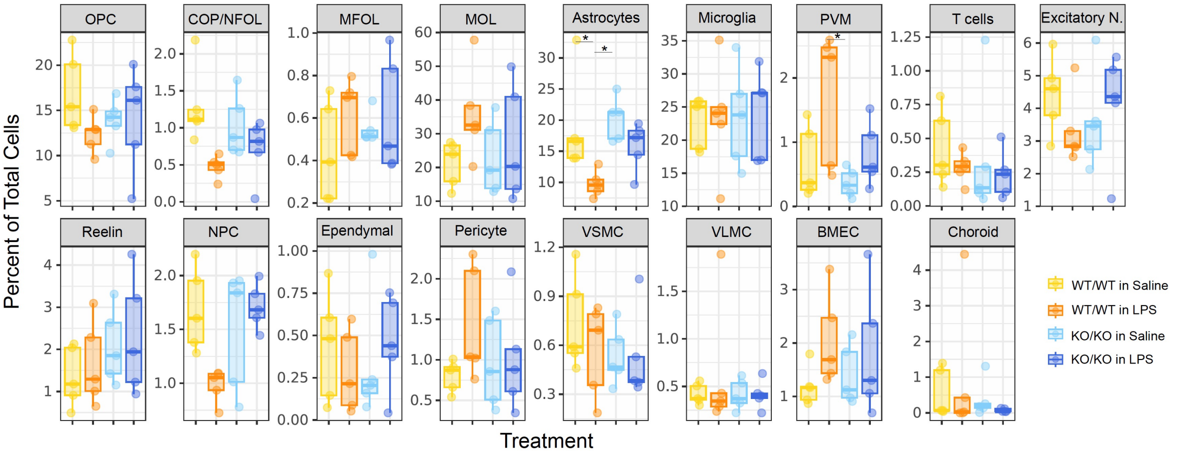


**Fig. S3 Changes in the proportions of major cell types by genotype and treatment effect.**

**(A)** Average expression of a canonical marker gene set for each major cell type identified in the dataset. **(B)** The percentage of cells in each cell type plotted as a percentage of the total number of cells within each sample (each dot is a sample). Differential abundance statistics * FDR < 0.05; ** FDR <0.01 and *** FDR< 0.001

Fig. S4

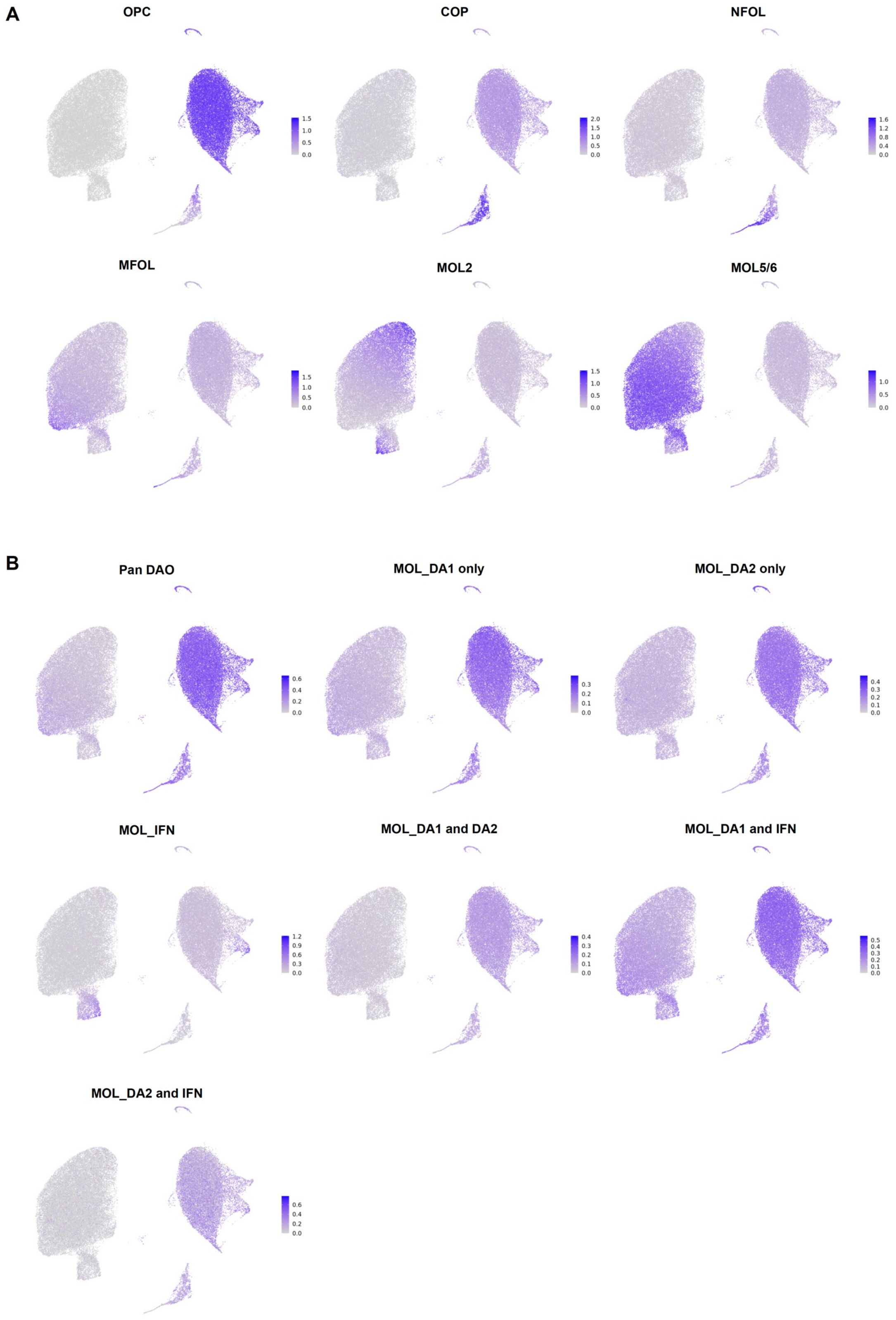


**Fig. S4 Mapping of DAO gene sets to OPC and oligodendrocyte subclusters**

**(A)** Average expression of a gene set that represents marker genes for each baseline OL lineage subtype. **(B)** Average expression of a gene set that represents marker genes for each disease-associated oligodendrocyte (DAO).
